# Colony entropy - Allocation of goods in ant colonies

**DOI:** 10.1101/572032

**Authors:** Efrat Greenwald, Jean-Pierre Eckmann, Ofer Feinerman

## Abstract

Allocation of goods is a key feature in defining the connection between the individual and the collective scale in any society. Both the process by which goods are to be distributed, and the resulting allocation to the members of the society may affect the success of the population as a whole. One of the most striking natural examples of a highly successful cooperative society is the ant colony which often acts as a single superorganism. In particular, each individual within the ant colony has a “communal stomach” which is used to store and share food with the other colony members by mouth to mouth feeding. Sharing food between communal stomachs allows the colony as a whole to get its food requirements and, more so, allows each individual within the colony to reach its nutritional intake target. The vast majority of colony members do not forage independently but obtain their food through secondary interactions in which food is exchanged between individuals. The global effect of this exchange is not well understood. To gain better understanding into this process we used fluorescence imaging to measure how the collected food is distributed and homogenized within a *Camponotus sanctus* ant colony. Using entropic measures to quantify food-blending, we show that while collected food flows into all parts of the colony it homogenizes only partly. We show that mixing is controlled by the ants’ interaction rule in which only a fraction of the maximal potential is actually transferred. This rule leads to a robust blending process: *i.e.*, neither the exact food volume that is transferred, nor the interaction schedule are essential to generate the global outcome. Finally, we show how the ants’ interaction rules may optimize a trade-off between fast dissemination and efficient mixing.

**Author summary:** We study how food is distributed in colonies of ants. Food collected by a small fraction of ants is distributed throughout the colony through a series of mouth-to-mouth interactions.

An interesting interplay exists between food dissemination and food mixing within the colony. High levels of dissemination are important as they ensure that any food type is available to any ant. On the other hand, high dissemination induces mixing and this reduces the required variety of nutritional choices within the colony.

Tracking fluorescent-labelled food and interpreting the results within concepts of information theory, we show that food collected by each forager reaches almost every ant in the colony. Nonetheless, it is not homogenized across workers, resulting in a limited level of mixing.

We suggest that the difference in food mixture held by each individuals can provide ants the potential to control their nutritional intake by interacting with different partners.

## Introduction

Food sharing in social insects is a compelling example of cooperation within a population [1–7]. Ants and bees can store a considerable amount of liquids in a pre-digestion storage organ called the ‘crop’ [8–10]. The stored food can later be regurgitated and passed on to others by mouth-to-mouth feeding (oral trophallaxis) [10–12]. Trophallaxis is a principal mechanism of food-transfer between individuals and therefore, the crop is often referred to as a “social stomach” [8].

When food is exchanged through trophallaxis, it is stored within the crop of the recipient workers and mixed with the rest of food in the crop [13–17]. Food blending is therefore an important factor in any process mediated by trophallaxis: from nutrient transfer and the maintenance of gestalt odor to hormonal regulation and information sharing [8, 13, 18, 19]. The extent to which food is blended in the colony has only been partially addressed before [3, 14, 20–22] and is still an open question.

Food blending is especially interesting in light of the fact that most colony members do not leave the nest [5, 14, 16, 23, 24], and all food is brought in by a a small fraction of workers called the foragers [16, 25]. The interplay between food-supplies brought in by different foragers can be expected to have an important role in the nutritional regulation of the colony. Social insect colonies have a documented ability to tightly regulate both the global nutritional intake [15, 21] and the dissemination of food to various sub-populations (such as nurses, larvae and brood) which may have different nutritional needs [5, 14, 16, 23]. The nature of this regulation is, however, not well understood.

Trophallactic food exchange requires physical contact between ants. The dissemination process is therefore conveniently described by a time ordered network, in which ants are the nodes and the food transfers are the (directed) edges. The topology of this network provides the underlying infrastructure of the food-sharing process [17, 26–28]. In the study of social insects and other real-world networks, the topology of the network can frequently be traced while the details of particular interactions are concealed [29, 30]. Indeed, previous studies that traced individuals in a colony have mainly focused either on the network topology [26, 28] or on coarse grained descriptions of food dissemination [1, 16, 22, 31]. In this study we use single ant identification and fluorescently-labeled food (Fig. 1a) to measure not only the interaction network but also the flow of food over this network.

**Fig. 1.**
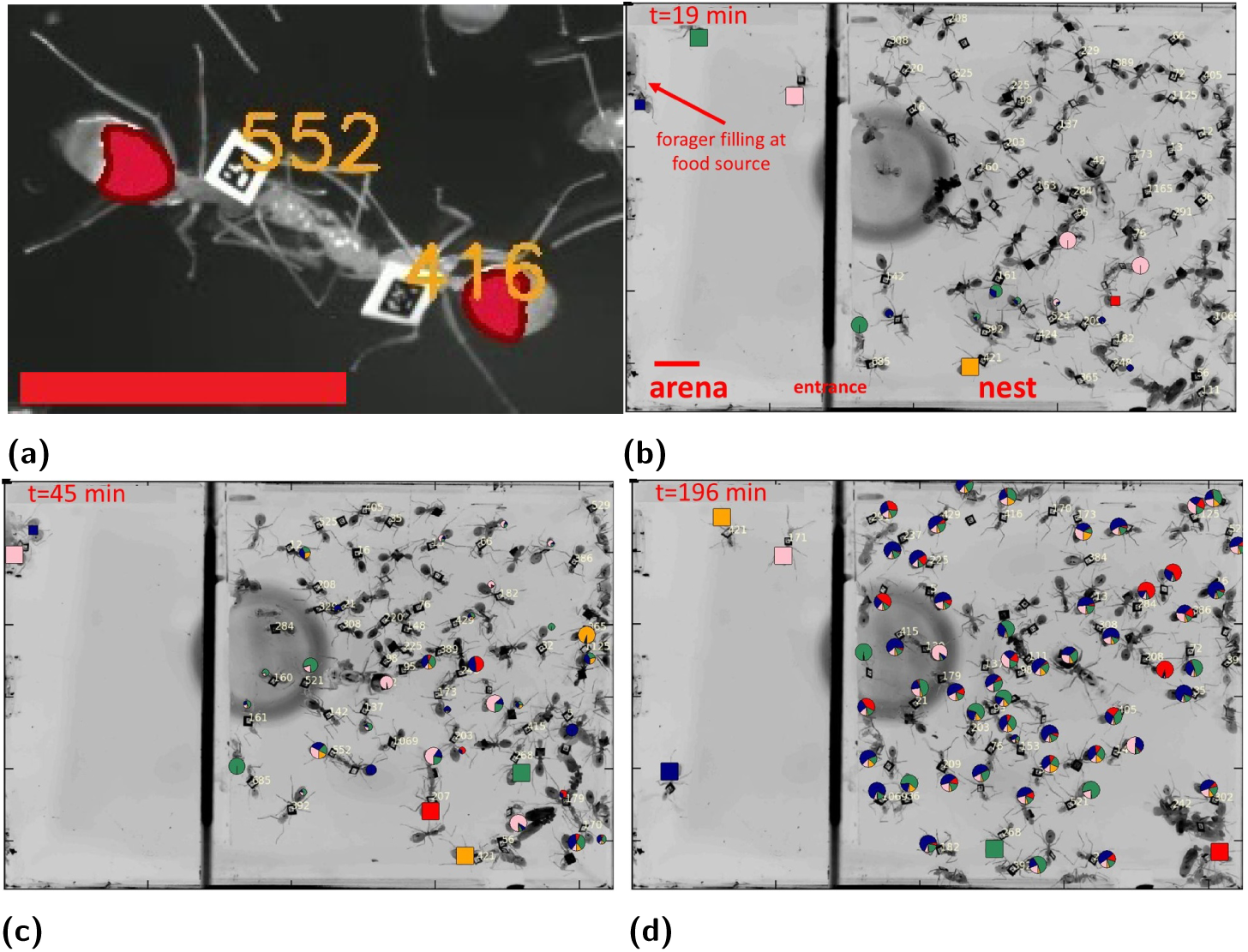
Experimental procedure. **a)** Two tagged workers engaged in trophallaxis. The identity of ants (orange numbers) was determined using Bugtag barcodes. The volume of food in the ants’ crop is measured using fluorescence imaging and overlaid in red. **b-d)** Food distribution across the colony and at different stages of the experiment. Markers (round: non-forager, square: forager) overlaid on ants depict their crop contents. Marker size is proportional to the food load held by each ant: *P*_*a*_ (small markers were set to a minimal size for clarity). Color division in markers of all ants depicts the computationally derived proportions of food in their crops according to the forager that first collected it (‘food-types’): (*P* (*f* |*A* = *a*)). Scale bar is 1cm. See also supplemental movie “Food dissemination in ant colony”.

The flow of food is limited by capacity: As the crop of ants is of finite size, this imposes a constraint on the amount of food that can be transferred in an interaction. This physical constraint limits the rate of mixing as ants become more and more full. Therefore, a potential trade-off between fast rate of food accumulation and well mixed outcome is expected.

The main objective of our study is using single ant measurement techniques to quantify how food blends as it is being disseminated across an ant colony. To this end, we use Shannon entropy to quantify the quality of mixing in ant’s crop. The Shannon entropy provides a single quantity that reflects the relative abundances of multiple constituents [32] and therefore sets a common scale by which food homogenization can be evaluated from our empirical data. Using our detailed measurements we characterize the interaction network and the rules by which food flows across this network. We then use hybrid simulations to identify which of these characteristics function as regulators of food mixing, and which might play a lesser role. Finally, we employ a theoretical model to study the trade-offs between food dissemination and nutritional homogenization.

## Results

### Food dissemination

We studied food (sucrose solution [80*g/l*]) dissemination in *Camponotus sanctus* ant colonies residing in an artificial, single chamber nest and following famine relief (see Materials and Methods, Experimental Setup). The dissemination process begins when the foragers, a small subgroup of the ants which we label *ℱ* = {1, 2, …, *N*_foragers_ *≡* |*ℱ*|}, return to the nest with liquid food loaded at the food source. Back in the nest, the foragers transfer the food to the non-forager population, *𝒜* = {*N*_foragers_ + 1, *N*_foragers_ + 2, …, *N*_ants_}, via trophallactic interactions (Fig. 1a). As food accumulates in the colony (Fig. 1b-d) it also flows between non-forager ants as they interact among themselves [17, 33].

The amount of food held in the crop of each ant as well as the amount of food passed per interaction were directly measured by combining single ant tracking with imaging of fluorescently labeled (Rhodamine B [0.08*g/l*]) food (Fig. 1a) [17]. We designate the total amount of food in the crop of a non-forager ant *a* at time *t* by *n*_*a*_(*t*) and the total amount of food held by all non-forager ants by *Z*(*t*) = Σ_*a∈𝒜*_*n*_*a*_(*t*). During the course of an experiment, the total amount of food held by the colony grows until it reaches saturation (Fig. S1e)^1^ [1, 33]. The fraction of the total food held by ant *a* by *P*_*a*_(*t*) = *n*_*a*_(*t*)*/Z*(*t*) is not uniform across colony members (Figs.2a, S1) and is restricted by variable physiological properties such as crop capacity.

As a first step towards quantifying food mixing in the ant colony we took a forager-centric approach. The idea is to track how food brought in by each forager spreads across the colony (Figs. 1b-d, 2a-b, S1c) and the degree to which these food flows may overlap and mix. Since our experiments included a single food source we implemented this approach using a computational procedure in which we define the type of each ‘food droplet’ by the index of the forager, *f ∈ℱ*, that had initially collected it at the food source (see ‘Food tracking’, Methods). This entails that the number of ‘food types’ in the system is taken to be equal to the number of foragers. Using the assumption that mixing of food inside the crop of an individual ant is extremely rapid when compared to the rate at which food is transferred between ants, we then tracked the trajectories of labeled food droplets as they flow through the colony (see ‘Food tracking’, Methods). This procedure allowed us to define the empirically measured probability, *P*_*a*_(*t*), described above (*P*_*a*_(*t*) can be viewed as the probability that a randomly chosen ‘food-droplet’ is found within the crop of ant *a*) and consider the inferred joint probability *P*_*f,a*_(*t*) = *n*_*f,a*_(*t*)*/Z*(*t*), which represents the probability that food, originally collected by forager *f*, is located in the crop of ant *a* at time *t*.

To quantify the degree to which different foragers contributed to the total foraging effort we calculate the total amount of food of type *f* that has accumulated in the colony up to time *t* as 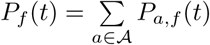. This probability function may be associated with an entropy, which we refer to as the *types-entropy* (*H*_types_), and which quantifies the relative abundance of the different food types (for all entropy definitions refer to SI, ‘Mathematical Framework’ and Table S2). It is defined by:

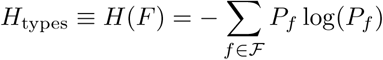

(we suppress the explicit notation of time from here onward). Our measurements show that *H*_types_ increases as a function of time (Fig. 2c) and quickly approaches the upper bound of log(|*ℱ*|). This upper bound can only be saturated if all foragers bring in equal amounts of food. As discussed below, *H*_types_ sets a limit on the total level of mixing in the colony.

**Fig. 2.**
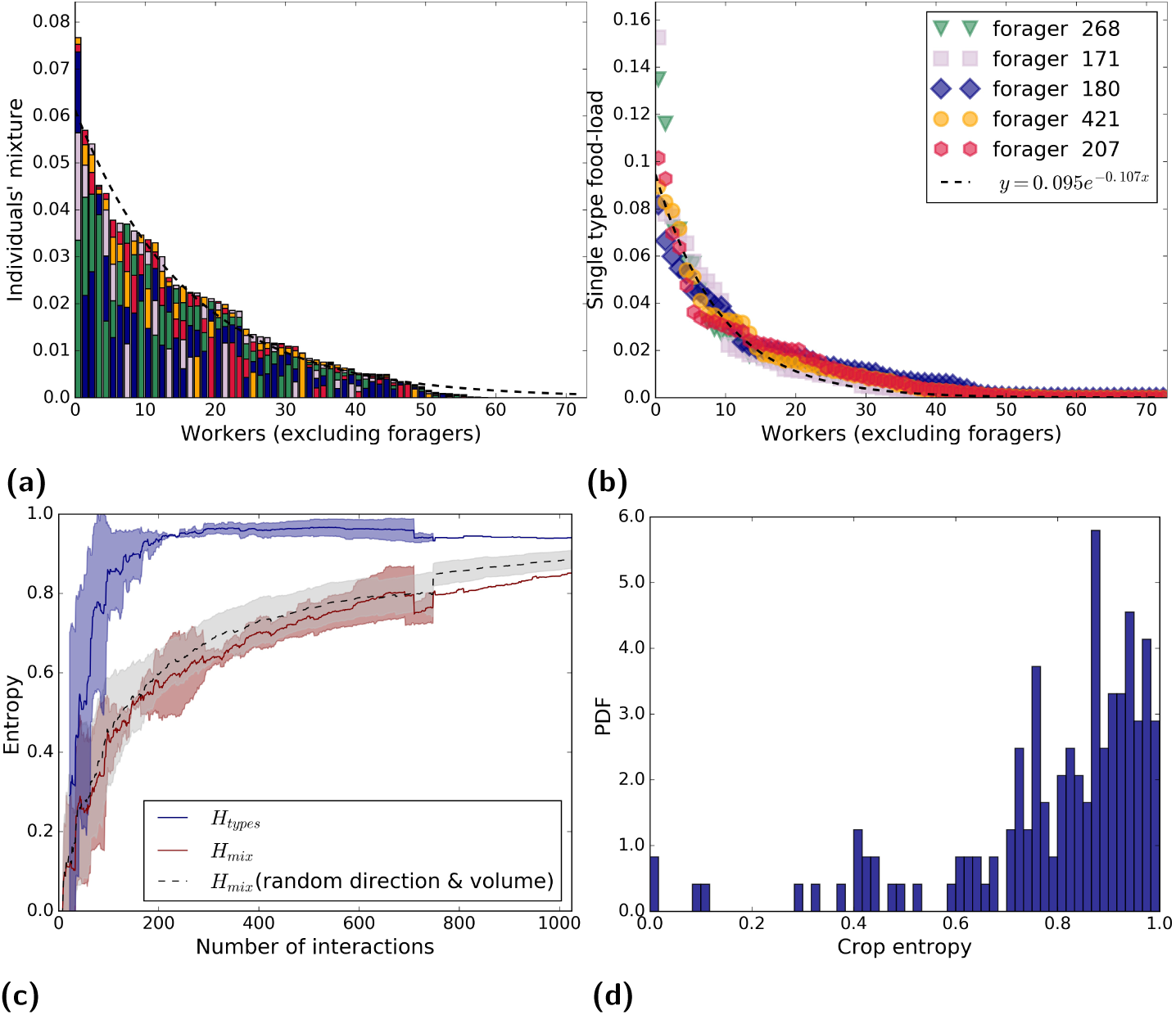
Food distribution. **(a)** The amount of food held by each non-forager ant *a* at the end of the experiment, *P* (*a*), colored by its forager origin *P* (*f* |*A* = *a*) (color code matches that of panel b, vertically ordered by amounts received). Ants are ordered by the amount of food in their crop, and the dashed line is an exponential fit *y* = *ae*^*-bx*^,*a* = 0.061 ± 0.002, *b* = 0.062 ± 0.003, *R*^2^ = 0.96. **(b)** The extent to which food from each forager *f* (color code as in panel a) was distributed among non-forager ants *a*: *P* (*a* | *F* = *f*). Recipient ants are ordered (per forager *f*) by amount received. Dashed curve is an exponential fit *y* = *ae*^*-bx*^, *a* = 0.095 ± 0.0013, *b* = 0.1 ± 0.002, *R*^2^ = 0.97. For colonies B and C see Fig. S2. **(c)** Mixing entropies as a function of the number of trophallactic interactions at the end of the experiment. Entropies are normalized by log(|*ℱ*|) to allow for data averaging over the three experiments. Lines are the mean over three experiments while shaded areas designate standard deviations. Depicted are the empirical entropy associated with the different proportions of food as brought in by each forager *H*_types_ (blue), the empirical mixing entropy over all non-forager ants *H*_mix_, (red) and the mixing entropy for hybrid simulations of randomized interaction volumes simulated over the empirical interaction schedule (*N* = 30, shaded area depicts standard deviation of the outcomes). Discontinuities are a consequence of the variable number of interactions among the three experiments. **(d)** A histogram of normalized individual mixing entropies of all non-forager ants, 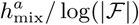, at the end of the experiments (all three experiments, *N* = 203 ants).

The degree to which food brought in by a single forager, *f*, spreads across the colony can be quantified by the conditional distribution *P* (*a* | *F* = *f*) = *P*_*f,a*_*/P*_*f*_. We found that the food initially collected by each and every forager reaches, practically, all members of the colony (Fig. 2b). This degree of dissemination dictates overlapping food flows such that the crops of non-forager ants hold a mixture of food of several types (Fig. 1b-d).

### Food mixing

Mixing was assessed by tracking the differently labeled food droplets as they flow, via the trophallactic network, from ant to ant. The conditional distribution *P* (*f* | *A* = *a*) = *P*_*f,a*_*/P*_*a*_ signifies the mixture of food-types in the crop of a specific ant *a* (Fig. 2a). Since each non-forager ant receives its load from multiple interactions with both foragers and non-foragers [17, 28] the food composition in her crop, *P* (*f* | *A* = *a*), contains a mixture of differently labeled ‘droplets’.

The level of blending in the crop of each individual ant, *a*, can be defined by the *crop entropy*:

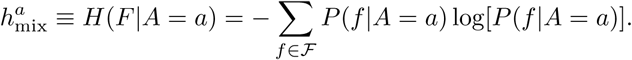

The range of individual crop entropy, 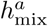, is [0, log(|*ℱ*|)] where zero crop entropy indicates that all food in the ants crop originates in a single forager while log(|*ℱ*|) indicates that food in the crop is equally divided among all possible food types. We find that the average mixing entropy (Fig. 2d) takes an intermediate value of 0.79 of the maximal possible mixing. While the actual components that mix to create the crop of each ant vary greatly (Fig. 2a) we find that the degree of mixing is actually quite uniform across the colony (standard deviation of 0.2 · log(|*ℱ*|), Fig. 2d).

Mixing within the entire colony, as a whole, can be quantified by the conditional entropy, *H*(*F* |*A*). This global *mixing entropy* is defined as the average over individual crop entropies, 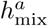, where each ant is weighted by its relative load, *P*_*a*_ [32]:

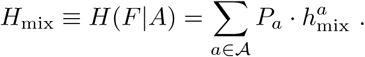

Mixing entropy is bounded from below by zero, a value which signifies no mixing: the food in the crop of any ant originates from a single forager only. An upper bound on mixing is obtained by the general rule *H*(*F* | *A*) ≤ *H*(*F*) (conditioning reduces entropy [32]) which, in our notation, translates into the fact that the mixing entropy is smaller or equal to the types entropy (*H*_mix_ ≤ *H*_types_). Equality signifies perfect blending and occurs only when all ants have identical crop-load compositions that exactly match the concentration-distribution of food types across the entire colony.

We find that as the number of interactions grows so does the mixing entropy, *H*_mix_ (Fig. 2c). However, while the crop composition of a typical ant contains food that originated from each of the foragers, the relative proportions of these food types differ from ant to ant and do not match the proportions of food as brought in by the foragers (Fig. 2a). In other words, even though the types entropy (*H*_types_) does approach the maximal bound of log(|*ℱ*|), the mixing entropy (*H*_mix_) is lower during the entire course of the experiment and reaches *H*_mix_*/H*_types_ = 0.8 ± 0.02 (mean ±std over three experiments) at the end of the experiments (Fig. 2c). If the mixing entropy does eventually reach the upper bound of the types entropy the time for this to occur is very long.

To discern the causes of these intermediate mixing levels we focus next on the underlying dynamics of food exchange. In the following sections, we characterize the pairwise interactions via which food spreads through the colony and study their implications on mixing.

### The process of food transfer

The flow of food across the colony can be described by focusing on two processes: 1) The *interaction network* which is the time-ordered depiction of the pairs of ants engage in trophallaxis. 2) *Interaction volume* which depicts food exchanged during an interaction in terms of both the direction and the volume. Next, we briefly characterize these two components.

#### Interaction network

Quantitative characterizations of temporal networks are difficult [35, 36], and in this section, we characterize network connectivity by studying the static graph which includes all interactions. In particular, we are interested in testing whether network connectivity (or its absence) may limit mixing. More accurate descriptions that take into account the temporal structure of the network will be discussed in the next section.

The *modularity* of a partition of a network into communities is defined as the fraction of the edges that fall within communities minus the expected fraction if edges were randomly distributed [37]. We used the ‘greedy modularity communities’ algorithm (‘Networkx’ Package for Python [34]) to search for a partition of the undirected trophallactic network which maximizes modularity. We found (Fig. 3a) that maximal modularity occurs for a partition into 3 - 5 communities, a number that is similar to previous estimates for this species [38]. However, these maximally modular divisions display low modularity (0.16 ± 0.024) in which the ratio of intra/inter edges tend to 1 (1.13 ± 0.3). We conclude that the food dissemination, most likely, is not hindered by missing links between communities.

**Fig. 3.**
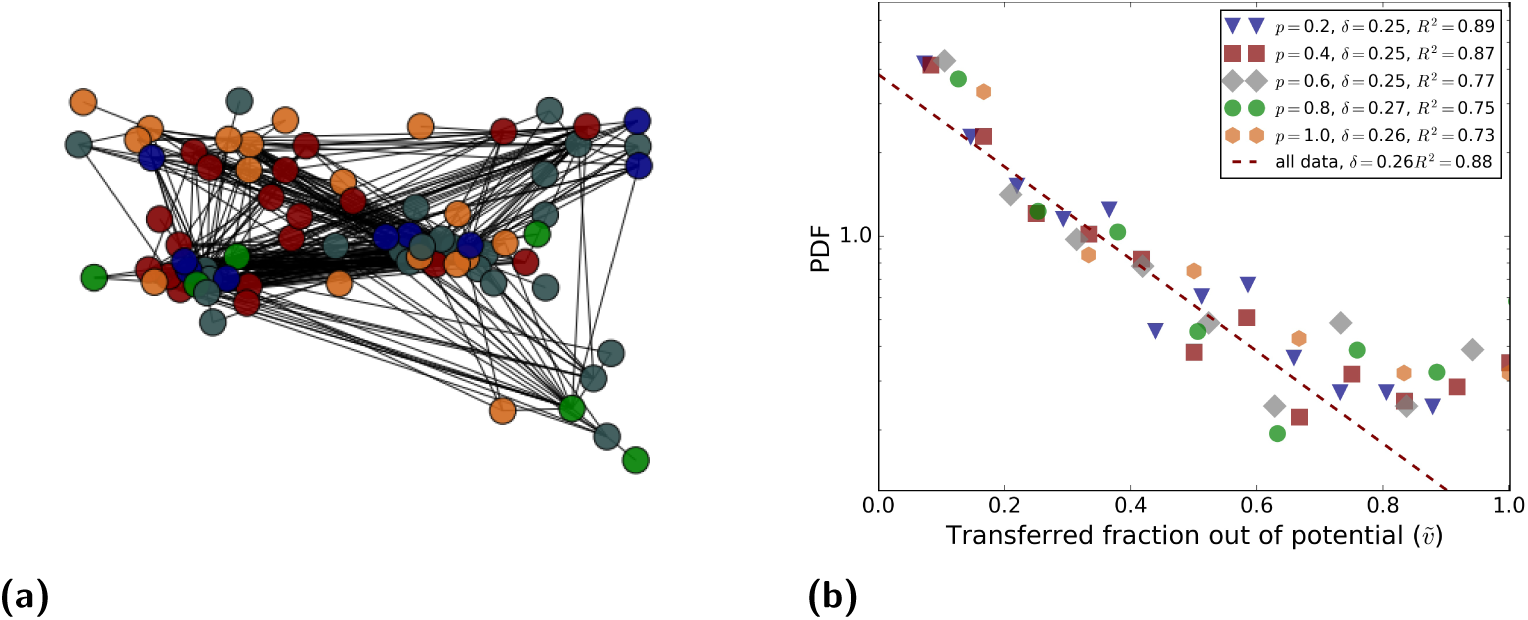
Pairwise interactions.**(a)** Visualization of the undirected trophallactic network (Colony A). Ants are represented by vertices and interactions are edges using the spring embedded layout from Networkx [34]. Nodes are colored according to maximally modular communities which, nevertheless, display low modularity with 191 intra-community and 223 inter-community links (Details: Number of communities=5, transitivity=0.38, modularity=0.185, quality performance=0.76). For colonies B and C, see Fig. S3. **(b)** Probability density as a function of the transfer ratio. Different colors relate to the maximal potential interaction *p* = *d*(1 − *r*) where *p ∈* [0, 1] is the volume potential as determined by *d, r ∈* [0, 1] donor’s and recipient’s crop load expressed as a fraction of the capacity of each ant. The distribution of interaction volumes per each potential are near-exponential functions: Fits to the exponential function 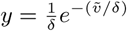 are provided in the legend (for different *p* values and for the entire data set (dashed line)). Data depict interactions from all three experiments (*N* = 2141 interactions).

#### Interaction direction and volume

When a forager ant interacts with a non-forager she typically acts as the donor (76 ± 16%, *N* = 713 where the error signifies interactions where direction could not be specified (Fig. S4a). When a forager was the recipient, the volume transferred was negligible (0.003 ± 0.008 mean ±std. *N* = 240) in comparison to all other cases (0.28 ±0.14 mean ±std. *N* = 1901).

To check for a possible choice of directionality in interactions between non-foragers we calculated the probability that a fuller ant (as a fraction of her own capacity) passes food to the emptier one. Within the limitations of measurements we find no such effect on the direction of transfer (56 ± 17%, *N* = 1357, error bars as above, Figs S4a,S4d). These results do not change when considering absolute amounts of the ants’ food-load rather than the fraction filled (Fig. S4b).

We next focus on *interaction volume*. As the crop of the ants is of finite size this limits the volume an ant can take. But the transfer is also limited by what the donor can give. Thus, we define the maximal transferable volume, *v*_max_, as the minimum between food in the donor’s crop and the free space in the receiver’s crop. The interesting finding is that, on average, the actual interaction volume, *v*, is no more than 0.26 ± 0.1, of this maximal potential regardless of how full or empty it and the donor-ant are (Fig. 3b). Furthermore, the distribution of the fraction 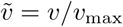 resembles an exponential:

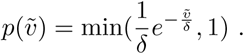

The fact that this *food-transfer rule* acts on 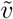 rather than on *v* itself suggests that, during interactions, ants control fractions of volume rather than absolute amounts [33]. Taking *δ* = 0.26 in the food-transfer-rule provides a good description of both the case in which the donor is a forager and the case where it is a non-forager (Fig. S4c, p-value *p* = 0.98, as computed by the Kolmogorov-Smirnov statistics on a set that includes only forager to non-forager interactions versus a set that includes only interactions between two non-foragers). It was previously suggested that, in interactions between foragers and non-foragers, it is the recipient ant that controls interaction volume [33]. The fact that the interaction volume rule does not depend on the identity of the donor is therefore consistent with an assumption that the recipient ant is not aware of this identity.

### Limits on macroscopic mixing

Different aspects of the trophallactic interaction may limit food mixing in different ways. One way in which mixing levels may be reduced stems from the details of the interaction rule. As an extreme example: if the crop capacity of all ants was about equal and in any trophallactic event ants would transfer as much food as possible this would lead to pure food loads that are simply relayed between the ants and therefore minimal mixing. Decreased mixing may also be the result an interaction network which is topologically segregated into several disjoint communities with limited food flows between them (as reported for other ant species [4]). In this section, we describe hybrid simulations, which preserve some of the empirically measured data while replacing others by simulated values (for details see SI, ‘Simulations’), to separately examine the effects of the different aspects of the interaction details on overall mixing.

#### Precise interaction volumes

Food mixing within the colony can be regulated by communication and controlled interaction volumes. For example, if interacting ants can sense that they hold very different crop compositions and react by reducing the trophallactic volume, this can limit mixing on the collective scale.

We have shown that the statistics of the trophallactic interaction volumes resembles an exponential distribution (Fig. 3b). In line with the above reasoning, this distribution might stem from precisely controlled interactions governed by internal parameters which we did not measure. However, an exponential distribution may also be the result of a random process where Poisson-like dynamics govern the termination of an interaction. A similar ambiguity holds for our measurements regarding the directionality of the interaction. As external observers, we have no way of telling random from non-random in this case. Nevertheless, we can assess the effect of possible randomness on the process of mixing.

To this end, we ran hybrid simulations in which ants interact according to the empirically measured network but in which interaction directions are chosen uniformly at random and interaction volumes are stochastically generated according to the empirical exponential food-transfer rule distribution (Fig. 3b). We find that these hybrid simulations exhibit limited mixing levels that are similar to the measured ones (Fig. 2c). In other words, the dynamics of food mixing does not suggest that the interacting ants use intricate communication, controlled directionality, or accurate interaction volumes.

#### Maximally mixing interaction rule

The clustering analysis presented above suggests that reduced levels of mixing are not a consequence of the network structure. However, this analysis was performed on a fixed interaction network that does not capture the temporal order at which interactions occurred.

To more accurately test whether interaction network properties limit mixing we ran hybrid simulations in which we kept the empirically measured interaction schedule including ant identities but replaced the measured interaction volumes by maximally mixing interactions: At each interaction, each ant gives half of her own crop to her trophallactic mate: Both ants leave the interaction with identical crop loads in terms of both volume and composition. Note that these interactions do not necessarily respect the empirically measured limited physical volume of each ants’ crop.

We find that the simulation curve of this ‘maximal-mixing’ rule exceeds the experimental data and leads to near maximal (*i.e., H*_types_) mixing levels (Fig. 4a). In other words, the connectivity and temporal structure of the trophallactic network can support maximal homogenization and are therefore not limiting factors on the mixing dynamics. This agrees with the observation that the static interaction network shows no clear community structure (Fig. 3a).

**Fig. 4.**
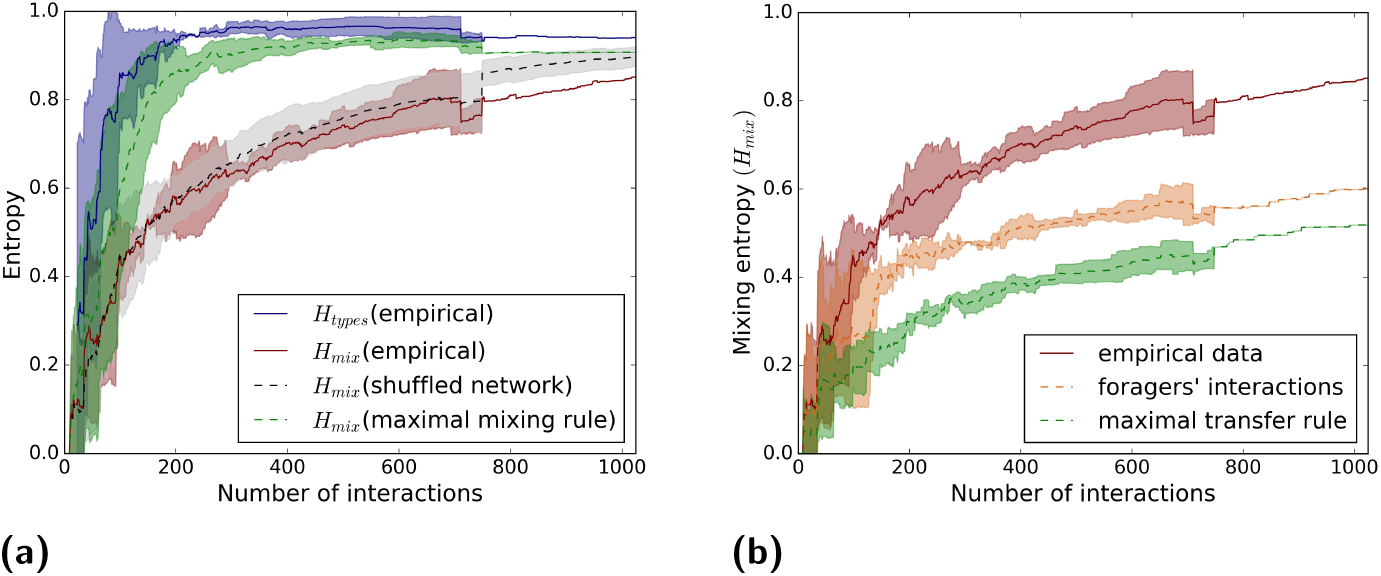
Mixing as a function of the number of interactions. Plots compare empirical data and hybrid simulations. Entropies are normalized by log(|*ℱ*|), solid lines show the empirical mean over the three experiments, dashed lines represent means over hybrid simulations. Shaded areas depict standard deviations. **(a)** The mixing entropy, *H*_mix_, in simulations with maximally mixing interactions applied over the empirically measured network (green curve) nearly saturates the empirically assessed upper bound *H*_types_ (blue curve). Mixing, *H*_mix_, in simulations where the empirically derived interaction rule is applied over maximally mixing interaction networks shows a limited rise which compares with empirical mixing rates (red curve). **(b)** Hybrid simulations of two extreme interaction rules preserving the empirically measured interaction schedule. The orange curve shows *H*_mix_ using only the transfers between foragers and non-foragers. The green curve depicts *H*_mix_ where every transfer is assumed to be at its maximal possible volume. These rules lead to mixing levels that are lower than those measured experimentally (red curve). Discontinuities in the plots are a consequence of the variable number of interactions among the three experiments.

#### Maximally mixing interaction network

To test whether the interaction rule limits mixing we replaced the empirical interaction network with randomly generated encounter patterns that are non-structured and do not inhibit mixing. These random networks were obtained by shuffling all individual identities in the measured interaction schedule table. This procedure allows us to preserve the number of interactions per individual while replacing the interaction network with one which is maximally mixing.

Since this shuffling process yields interactions that did not actually occur we used the empirically measured food-transfer-rule to simulate random interactions (in both directions and volume) over the simulated networks. As shown in the previous sections and in Fig. 2c, this replacement is not expected to have implications on the global mixing process.

We find that simulated mixing over shuffled networks did not show any statistically significant deviation from the empirically measured mixing process (Fig. 4a). We therefore conclude that the statistical properties of the ants’ interaction rules, which respect the physical capacity of the crop, will limit food mixing within the colony.

#### Extreme interaction rules

Our results thus far raise the interesting possibility that the value of *δ* is a regulator of mixing. To explore this direction, we start by looking at two extreme cases: *δ* = 0 (for non-forager donors) and *δ* ≫ 1 (for all ants).

We first ran hybrid simulations in which all forager to non-forager interactions were maintained at their empirical values (corresponding to *δ* = 0.26) while all interactions among non-foragers were omitted (corresponding to *δ* = 0). We find that this manipulation reduces mixing levels and that interactions between pairs of non-foragers indeed contribute to food mixing within the colony (Fig. 4b).

The interaction dictated by *δ* ≫ 1 is that of maximal transfer. Here, the empirically determined donor-ant was simulated to transfer as much as it could possibly pass given the trivial volume constraints. In other words, the amount passed was set to be the minimum of two volumes determined prior to the interaction: what the donor holds and the empty space in the receiver’s crop. Similar to the previous case, the maximal transfer rule shows a reduced level of mixing (Fig. 4b). Note that mixing levels do not go to zero. The reason is that the crop capacity is not equal across all the workers so when food is transferred it is also divided.

### Trade-off between mixing and accumulation rates

Finite crop size naturally impacts an ant’s ability to mix food. Mixture composition can significantly change only if an ant receives a large enough portion relative to her temporal load. Therefore, as ants become more satiated, their free storage space (*i.e.*, the difference between her capacity and her current load) becomes smaller and the ability to mix (the potential mixing rate) declines. Consequentially, a fast accumulation rate might interfere with the mixing process.

As implied from the empirical interaction rule, in a receiving interaction, an ant is provided with a random volume of food that follows an exponential distribution, with an average that is proportional to her free storage space. This means that on average, an ant receives food in a series of decreasing volumes with a parameter *δ*. The parameter *δ* can thus be expected to have opposite effects on the accumulation and mixing of the food: the larger the value of *δ* the higher the accumulation rate and the lower the mixing rate (and *vice versa*).

We used a simple model to explore the possible trade-offs between the rate at which food accumulates within the colony and the extent to which it is mixed. The model assumes that all ants have the same capacity (for simplicity), that foragers and non-foragers use the same 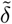 (in a deterministic version of the original food-transfer rule, see SI, ‘Simulations’) and that interactions occur randomly. Furthermore, for the purpose of the model, we defined the amount of food held by a forager at time *t* = 0 to equal the total amount of food she collects at the food source during the entire course of the experiment. This definition sets the amount of food across all colony members, *M*, as a quantity that is conserved over time. Considering the entire colony we now define the probability 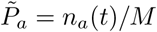 as the fraction of total amount of food held by any ant, forager or non-forager.

Using these definitions entails that at *t* = 0 all food is held by the foragers being, therefore, completely non-mixed while at later times, as food flows into the colony, it mixes within the crops of non-forager ants. This interplay between food accumulation and food mixing can be captured by considering the mixing entropy over all ants in the colony:

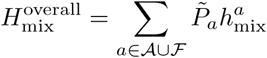

Note that since foragers receive almost no food from other workers (see above) we can approximate *P* (*f′*|*a* = *f*) *≈* 1 for *f′* = *f* and zero otherwise. This means that 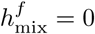 for *f ∈ℱ* and leads to a second representation of *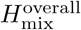* (see SI, ‘Trade-off model’):

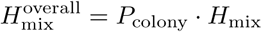

where 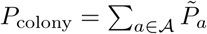 is the colony’s satiation level which starts off at 0 and saturates at 1 as food flows into the system [17]. This representation neatly separates the dissemination behavior into a component which quantifies the extent at which food is accumulated and a second component which quantifies the extent at which it is mixed.

We simulated an approximation to this model (see SI, ‘Simulations’) to study the relative effects of these terms as a function of the parameter 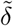. The interactions of the simulations approximate the empirical data by keeping the average interaction per ant and the ratio between forager to non-forager and non-forager to non-forager interactions. As may be expected, larger values of the parameter 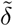 lead to larger transfer of food into the colony (*P*_colony_ indicated by the green line in Fig. 5a). However, due to the finite capacity of an ant’s crop, larger values of 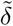 also hamper mixing among non-foragers (*H*_mix_ indicated by the blue line in Fig. 5a). The compromise between these two factors is captured by the product of these two, the total mixing entropy 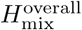 which exhibits a maximum for an intermediate value of *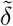*.

**Fig. 5.**
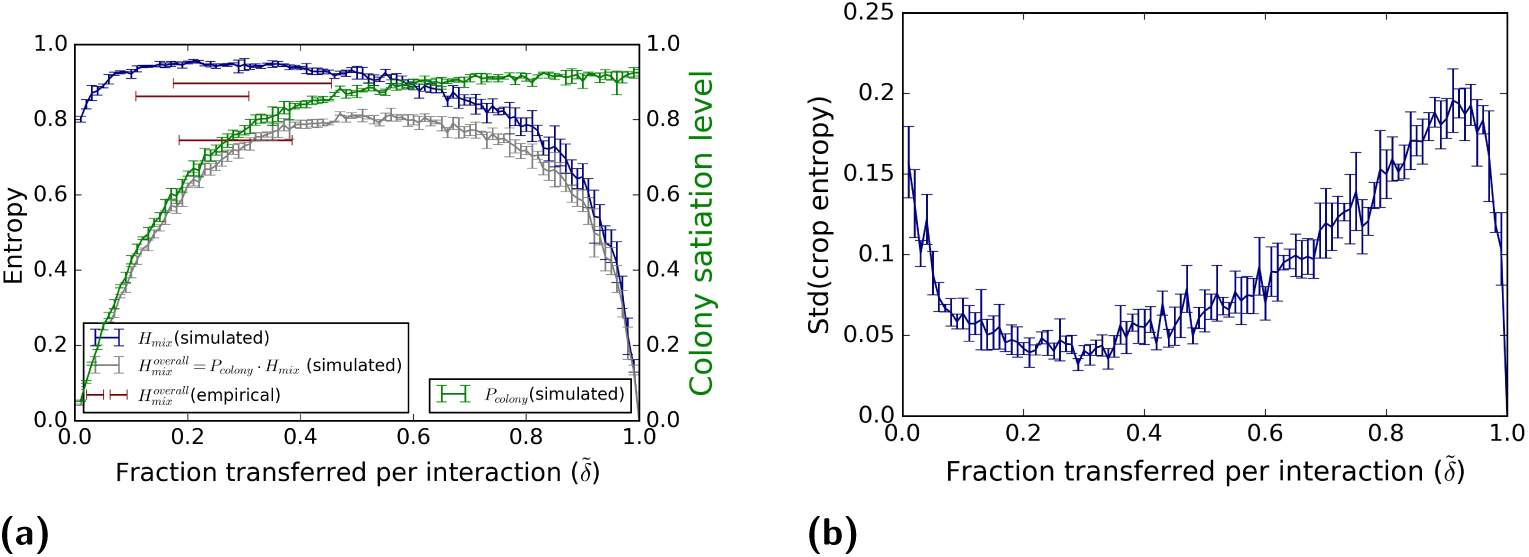
The 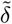 Model. **(a)** While the colony state (*P*_colony_, green) rises with the fraction 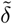 of transferred volume, mixing levels among non-foragers (*H*_mix_) decreases. The mixing levels over all ants in the colony (including the foragers) is the product of these two functions, 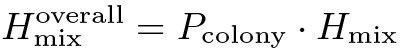 and displays a broad maximum which spans all non-extreme values of 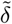. Curves can be compared to the empirically measured values (red bars) of the three experiments. **(b)** Standard deviation of individual mixing entropies 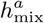 across all ants. Standard deviations were calculated for 30 model runs. The plot depicts the mean and standard deviation of this value.

These results demonstrate a robust process: as long 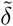 does not approach the extremes, both the mixing and the accumulation are comparable for a given number of interactions (Fig. 5a). Surprisingly, even though higher *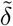* will result in a higher accumulation rate, the ants seem to function at smaller 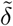 values (red bars in Fig. 5a). A potential benefit of smaller 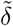 values is the maintenance of similar mixing levels across all ants in the colony (Fig. 5b). This stands in agreement with our empirical evaluation of the variance in mixing levels across the colony (Fig. 2d).

## Discussion

It is well known that social insects manage their nutrient resources on the collective level and also on finer scales because the colony channels foods with different nutritional composition to different sub-populations. In this paper, we put forward the idea that this intricate regulation relates to the interplay between food dissemination and food mixing within the colony. High levels of dissemination are important as they ensure that any food type is available to any ant. On the other hand, high dissemination induces mixing and this reduces the required variety of nutritional choices within the colony.

A main finding of this work is that, despite repeated trophallactic interactions between the ants, food in the colony does not become evenly mixed. Quantifying mixing using entropy measures we showed that, compared to what was theoretically possible, mixing is slow to rise and levels up at around 80% of the full mixing potential. The logarithms in the definition of entropy make the significance of this number difficult to assess. For intuition, in the case of only two food sources, the maximal mixing entropy (1 bit) corresponds to each crop holding equal parts of the food sources (1 : 1) while 80% of this (0.8 bits) corresponds to, a far from perfect, 3 : 1 partition of food sources. This imperfect mixing offers the possibility for ants to choose from a wide spectrum of nutritional compositions when the donors provide different blends. Such a choice can allow ants within the nest to reach their nutritional target using feeding schemes similar to those described by the geometrical framework for food foraging [39, 40].

We further explored the mechanisms that allow for intermediate levels of food blending. Using hybrid simulations, we found that the interaction network over which food flows does not pose any limits on mixing levels. Rather, it is the interaction rule employed by the ants that regulates the extent to which food blends. This is reminiscent of several examples in which cellular pathways with identical architecture can achieve starkly different regulatory behaviors depending on actual rate coefficients [41, 42]. Regulation by interaction rules rather than by meeting patterns is an intriguing possibility for social insects in which different collective functions often reside over very similar interaction networks [26]. For example, while proximity is required for both food sharing and disease transmission [43] different interaction rules may ensure that one of these is enhanced while the other is suppressed.

Quantifying a large number of trophallactic interactions, we directly measured the food-transfer rule (see also [33]) used by the ants. We stress several important aspects of this rule. First, the rule, naturally, respects the physical limits on crop size of the ants. Broadly speaking the food-transfer rule implies that a few interactions will be larger than most others. Therefore, to a great extent, the entropy of the food mixture within the crop of an individual is determined by a few voluminous events and does not reach the maximal limit (see SI, ‘Entropy by largest events’, Fig. S5). Second, we show that the interaction rule is most likely stochastic in nature and, therefore, does not entail any strong requirements on ant cognition or communication. Finally, the fact that in trophallactic interactions the recipients fill only partially (Fig. 3b) is in agreement with a model in which, similar to animals foraging in the environment, ants in the nest regulate their nutritional income by feeding off of multiple partners each with a different mixture of the available ‘food types’.

We explored the interplay between food dissemination and mixing using a simple model of food flow that is based on our empirical observations. We find that the intermediate levels of mixing, as measured, can viewed as a compromise between the requirements to quickly unload incoming food and the requirement to disseminate different food types to all parts of the colony. We show that this process is robust over a wide range of *δ* values and that the actual measured parameter ensures that all ants in the colony are equally well mixed (although each holds a different particular mixture).

Finally, we wish to highlight the limitations of this study. Due to current technological availability, this work was performed using a single food source labeled by a single dye. The ants may behave differently in terms of both interaction network and food-transfer-rule when several food sources with different nutritional values are available [4]. Further, our artificial setup contained a single chamber nest. More realistic, multi-chambered, nest structure may induce interaction networks that are more clustered than the one measured here. This may hold important consequences for nutrition dissemination. Last, is our choice to measure mixing by labeling food types by foragers. While arbitrary, this is a reasonable choice since, as we have shown, foragers are responsible for a large part of the mixing (Fig. 4b). Taking all these limitations into account we view our findings as a baseline to which future results in more complex situations may be compared to.

Overall, our finding that the interaction rule takes precedence over the interaction schedule manifests both the robustness of collective processes within the ant colony and the large extent to which individual behaviors may modulate global outcomes.

## Materials and methods

For a more comprehensive methods section please refer to the SI and [17, 33]. Our experiments were conducted on lab colonies of *Camponotus sanctus* which included 50-100 workers, reared from single queens that were collected during nuptial flights in Neve Shalom and Rehovot, Israel. Table S1 contains further details on each experimental colony.

### Experimental setup

The experimental setup consisted of an IR-sheltered artificial nest chamber (∼100 cm^2^), neighboring an open area which served as a yard. The setup was recorded by two cameras (details in [17]): the top camera images were used to extract ant identities, coordinates and orientations using the BugTag software (Robiotec). The bottom camera images were used to detect fluorescent-labelled, using the openCV library in Python. Combining the information from both images, we associated between the identity of an ant and her appropriate fluorescent image. Thus, for each experiment a database was obtained, which included for every frame the coordinates, orientation, and measured fluorescence (in arbitrary units of pixel intensity) of each identified ant.

### Food tracking

The experimental trophallactic network includes a time-ordered pairwise-interaction schedule, and the volume of liquids that one ant passed/received to (from) the other. Food is tracked from the moment it is acquired by a forger from the food source. We associate this volume (food ‘droplets’) with the forager’s barcode identity (‘type’), and continue tracing these droplets as they split between the ants according to the interaction schedule. To do this, we assume that in each interaction the receiver ant receives food volume with the same distribution of types of food droplets as in the crop the donor, *i.e.*, and the amount of food droplets of type s received by the recipient is equal to:

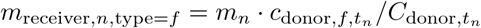

Here, *m*_*n*_ = total amount of food obtained in the event *n, m*_receiver,*n*,type=*f*_ is the amount of food the receiver ant received in interaction *n* that originally came from the forager *f*. Also, *t*_*n*_ is the beginning time of interaction *n* and 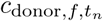 is the amount of food in the crop of the donor ant that originally came from forager *f* just before the interaction began. Finally, 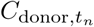 is the total amount of food possessed by the donor ant at time *t*_*n*_, *i.e.*, just before the interaction begun.

Doing this, we obtain an estimation of the number of food droplets of type *f* within the crop of ant a at any time t(*n*_*a,f*_ (*t*)), which we then use to calculate entropy (see SI, mathematical Framework).

## Author Contributions

E.G and O.F. designed the experiments. E.G. performed the experiments and conducted the analysis and simulations. E.G, O.F, and J-P.E. developed the analysis methods and wrote the paper.

## Declaration of Interests

The authors have no competing intersests.

## Acknowledgements

We would like to thank Lior Baltiansky, Elisha Moses, and Amos Korman for useful discussions. We would further like to thank Guy Han and Yuri Burnishev for technical help. O.F. was supported by the Israeli Science Foundation grant 833/15, and the European Research Council under the European Union’s Horizon 2020 research and innovation program (DBA-648032).

## Supporting information

### SI Methods

#### Study Species: ***Camponotus sanctus***

*Camponotus sanctus* are omnivorous ants that are presumed to naturally live in monogynous colonies of tens to hundreds of individuals (projecting from *Camponotus socius*, [44]), distributed from the near East to Iran and Afghanistan [45]. Workers of this species are relatively large (0.8-1.6 cm) and characterized by translucent gasters, rendering them suitable for both barcode labeling and crop imaging. Our experiments were conducted on lab colonies of 50-100 workers, reared from single queens that were collected during nuptial flights in Neve Shalom and Rehovot, Israel. Table S1 contains further details on each experimental colony.

**Table S1.** Experimental Colonies.

| Colony | # Ants | Starvation(weeks) | # Major Workers* | # Foragers |
| --- | --- | --- | --- | --- |
| A | 100 | 3 | 1 | 5 |
| B | 62 | 5 | 1 | 3 |
| C | 53 | 4 | 2 | 4 |

\* ‘Major Workers’ refers to the cast of workers in the colony which have distinctively larger body size.

#### Experimental setup

Fluorescent food imaging and 2D barcode identification (BugTag, Robiotec) were used to obtain a live visualization of the food flow through colonies of individually tagged ants. See [17] for a detailed description of the experimental setup. In short, an artificial nest was placed on a glass platform positioned between two cameras. A camera below the nest filmed through the platform, capturing the fluorescence emitted from the food inside the translucent ants. Meanwhile, a camera above the nest filmed through its infrared shelter, capturing the barcodes on the ants’ thoraxes, allowing identification of single ants inside the nest. Together, footages from both cameras enabled the association between each individual ant and her food load, throughout time and across trophallactic events. The two cameras were synchronously triggered at a fixed frame rate, (here 0.5 Hz., except for colony B which was recorded at 1 Hz.). We chose a temporal resolution that is sufficient to capture events of 2 seconds since shorter interactions barely involve food exchange [17].

#### Image processing

Top camera images were used to extract ant identities, coordinates and orientations using the BugTag software (Robiotec). Bottom camera images were used to detect fluorescence with a pixel intensity threshold, using the openCV library in Python. Gasters of fed ants appeared as bright “blobs” and thus passed the image threshold (for details, see [17]).

In order to associate between the identity of an ant and her appropriate blob, the image from the upper camera was transformed to align with the fluorescent image. Then, for each identified tag, a small area extended from the back of the tag toward the ant’s abdomen was crossed with the thresholded fluorescent image. If a blob intercepted this area, it was assigned to the tag’s identity.

Thus, for each experiment a database was obtained, which included for every frame the coordinates, orientation, and measured fluorescence (in arbitrary units of pixel intensity) of each identified ant.

#### Experiment protocol

Following a food-deprivation period of 3-5 weeks, ant colonies (queen, workers and brood) were manually barcoded and introduced to the experimental nest for an acclimatization period of at least 4 hours. The nest consisted of an IR-sheltered chamber (∼100 cm^2^), neighboring an open area which served as a yard. After the acclimatization period, the two cameras synchronously started to record. After 30 minutes, the fluorescent food (sucrose [80 g/l], Rhodamine B [0.08 g/l]) was introduced to the nest yard *ad libitum*, and the recording proceeded for at least 4 more hours - a duration sufficient for the colony to reach its desired food volume intake [33].

#### Foragers

Each experiment consisted of a few individuals who performed consistent foraging cycles between the food source and the nest. Those ants were considered as “foragers”. Some other individuals were occasionally observed at the food source but clearly did not display such foraging cycles. For our purposes they were not considered as foragers. These ants visited the food source no more than 4 times, while consistent foragers performed an average of 15.67 cycles and no less than 8. The data presented here is from the first return of a forager to the nest from the food source until the end of the experiment [33].

#### Interaction identification and crop load estimation

Even though the fluorescence emitted from an ant’s crop is reasonably indicative of the food volume, it is a noisy measurement mainly due to her highly variable postures. Therefore, assuming that an ant’s crop content remains constant during the intervals between trophallactic events, we evaluated the temporal food load by 90^*th*^ percentile fluorescence measurement acquired in each such interval [17].

In order to precisely consider the relevant intervals for this estimation, the trophallactic interactions were manually identified from the video. Interactions were classified as trophallactic events whenever the mandibles of the participating ants came in contact and the mandibles of at least one of the ants were open. For forager ants, another situation in which their crop loads may change is when they directly feed from the food source. These feedings were also manually identified from the video, as times when a forager’s open mandibles touched the food source.

#### The volume of trophallactic event

For each trophallactic event we recorded two measurements: one of each participating ant. While the transparency factors (calculated for each ant relative to the others [17]) reduce discrepancy between these two measurements they do not eliminate them completely. Thus, the amount of food transferred in an interaction *n* is calculated by a weighted average corresponding to a maximal likelihood of normal distribution:

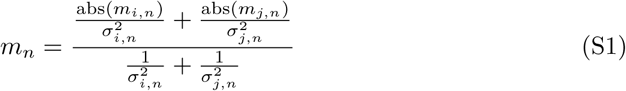

Where: *m*_*n*_ - the estimate amount transferred in interaction *n*.

*m*_*i,n*_ - is the amount transferred in interaction n according to ant 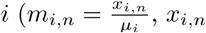 - difference in pixels value according to ant *i, µ*_*i*_ - transparency factor associated with ant *i* see, [17]) 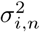 - measurement error (variance of the measurement points, standardized to the number of measurements in time intervals before and after the interaction) of *m*_*i,n*_.

#### Simulations

##### Maximal mixing

the simulation follows an experimental interaction schedule (*i.e.*, pairs, time-order and direction) while ‘equal sharing rule’ is applied. In this case, at each interaction, each ant gives half of her own crop to her trophallactic mate. This is the best mixing scenario that can be achieved in a single interaction since both ants leave the interaction with identical crop loads. This interaction rule further translates to maximal mixing on the global level. Specifically, in each step of the simulation the maximal exchange rule is applied between the empirically determined interaction partners and their crop loads are accordingly updated (see above ‘food tracking’, methods). Finally, *H*_mix_ is calculated for the resulting food distributions.

##### Maximal transfer

the simulation follows an experimental interaction while ‘maximal transfer rule’ is applied. In this case, at each interaction, the experimental donor-ant gives the maximal amount that could have passed *i.e.*, min(donor’s food load, recipient’s capacity - recipient’s food load) and *H*_mix_ is calculated as described above.

##### Foragers only

the simulation follows an experimental interaction schedule while only interactions that include foragers are taken into account. The transferred amount is taken from the experimental data and *H*_mix_ is calculated as described above.

##### Random rule

This simulation follows the time-order of pair-trophallaxis. In trophallaxis events between foragers and non-foragers the forager is assumed to be the donor, and in all other cases the donor is chosen randomly and with equal probably between the experimental pair. The transferred volume, *v* = *r · v*_*p*_, is determined by the multiplication of a random factor *r* (*r* = min(*x*, 1) where 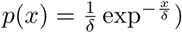) and the potential volume *v*_*p*_ (*v*_*p*_ =min(donor’s food load, recipient’s capacity - recipient’s food load)). *H*_mix_ is calculated as described above.

##### Shuffle simulation

We separately shuffle the donor time ordered-list and the recipients’ list and create a new trophallaxis-network by combing these lists together (*i.e.*, the identity of the trophallaxis-mates, and hence the direction of the trophallaxis events are randomly changed). The transferred volume and *H*_mix_ are calculated as described for the random-rule above.

##### Trade-off simulation

. This simulation approximates food flows in the trade-off model as described in the main text. In this simulation all ants are assumed to have the same capacity. We begin each run (in total 30 runs per given 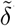) by creating a random trophallaxis network that includes 60 ants in total, 4 foragers and 1000 interactions We use a ratio of 3:1 between interaction that include foragers and ones that do not. These parameters of the simulations were chosen to approximate the empirical data in terms of number of ants, number of foragers, the average interaction per ant and the ratio between forager to non-forager and non-forager to non-forager interactions. At time zero, all ants are empty, except the foragers who are always completely full. The simulation follows the randomly generated network, and the transferred volume is taken to be a constant fraction, 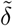, of the potential that could have passed (this is a deterministic approximation of the empirical random transfer rule described by the parameter *δ*). Here again, if a forager is involved in the trophallaxis she is taken as a donor, otherwise the donor is determined randomly. Each time step includes one trophallaxis event, in which the donor and transfer fraction 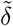 out of the potential the could pass, *n*_*a,f*_ (*t*) is reevaluated for trophallaxis mates. We use this data to calculate

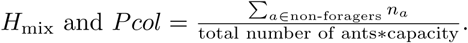

### Supplemental Text: Mathematical Framework

#### Colony Entropy

In our experiments, only one food source was provided: a sucrose solution. Therefore, to study the mixing, we labeled each ‘food droplet’ by a type, according to the forager that brought it(see Methods). Therefore, the number of ‘food types’ equals to the number of foragers in the experiment.

*ℱ* = {1, 2, *…, N*_foragers_ *≡* |*ℱ*|} - the set of foragers in the colony.

*𝒜* = {*N*_foragers_ + 1, *N*_foragers_ + 2, *…, N*_ants_}- the set of non-forager workers in the colony.

By tracking the ‘food droplets’ as they passed in trophallaxis we could calculate the joint probability, *P*_*a,f*_, that a particle of source *f* is in the crop of ant *a*:

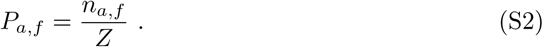

where,

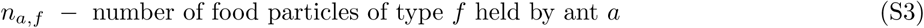

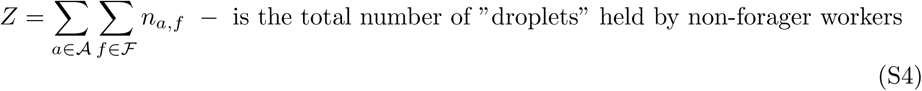

With this joint probability the total Entropy of the colony, *H*_colony_, is defined as the Shannon Entropy (SI Fig. S1a-S1b):

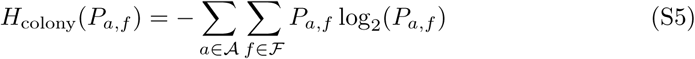

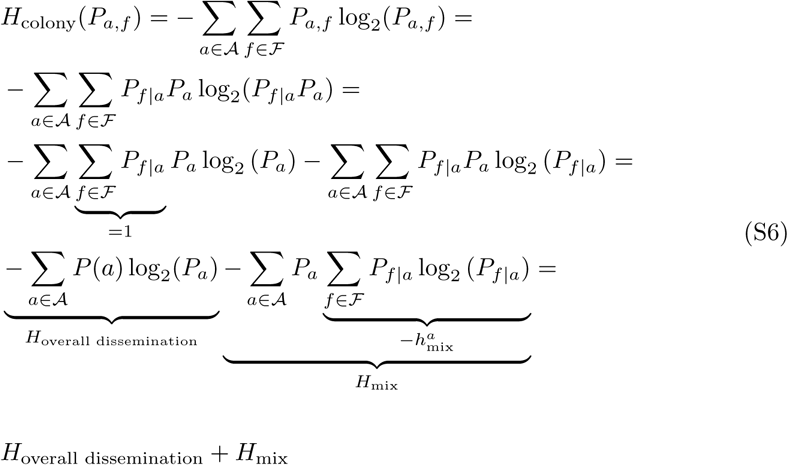

and the following identities were used:

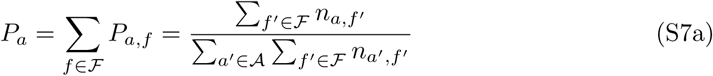

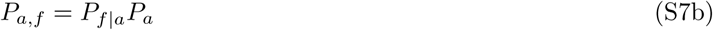

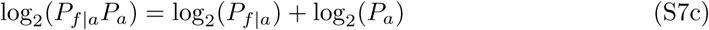

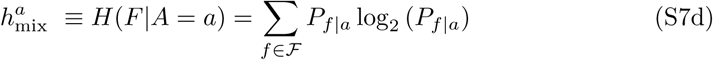

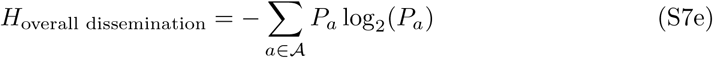

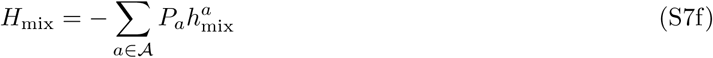

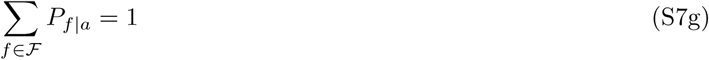

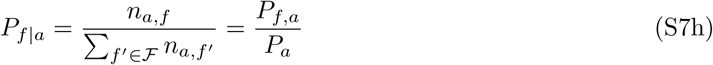

In a similar way, the entropy of the colony can also be divided into the two components, types and foragers’ dissemination (SI Fig. S1c-S1d):

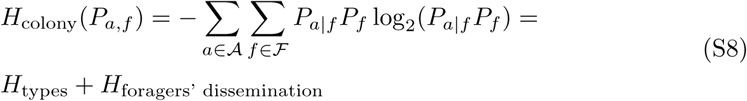

where:

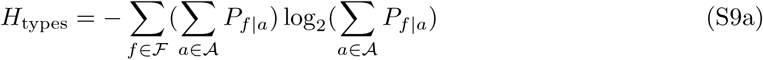

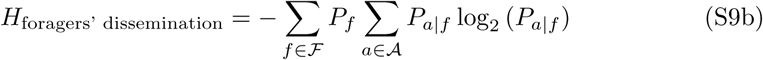

Each of these four conditional entropies may be related to a different aspect of the collective level of the food dissemination process:

1. *H*_overall dissemination_ - How uniform is the food divided across the workers of the colony (regardless the food type)?
2. *H*_mix_ - How well food is mixed across individuals?
3. *H*_types_ - The abundance of each food type
4. *H*_foragers’ dissemination_ - How uniform each food type is distributed across the ants?

For the dynamics of these different entropy terms please refer to figure S1.

#### Trade-off model

For the purpose of the model, we defined the amount of food held by a forager at time *t* = 0 to equal the total amount of food she collects at the food source during the entire course of the experiment. This definition sets the amount of food across all colony members, *M*, as a quantity that is conserved over time. Considering the entire colony we now define the probability 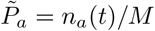 as the fraction of total amount of food held by any ant, forager or non-forager.

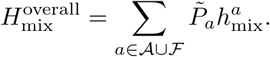

Note that since foragers receive almost no food from other workers we can approximate *P* (*f ′*|*a* = *f*) *≈* 1 for *f ′* = *f* and zero otherwise. This means that 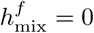 for *f ∈ℱ* and leads to a second representation of 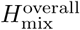:

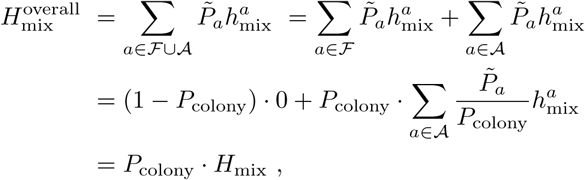

where 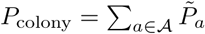 is the colony’s satiation level which starts off at 0 and saturates at 1 as food flows into the system [17]. This representation, therefore, neatly separates the dissemination behavior into a component which quantifies the extent at which food is accumulated and a second component which quantifies the extent at which it is mixed.

**Table S2.**
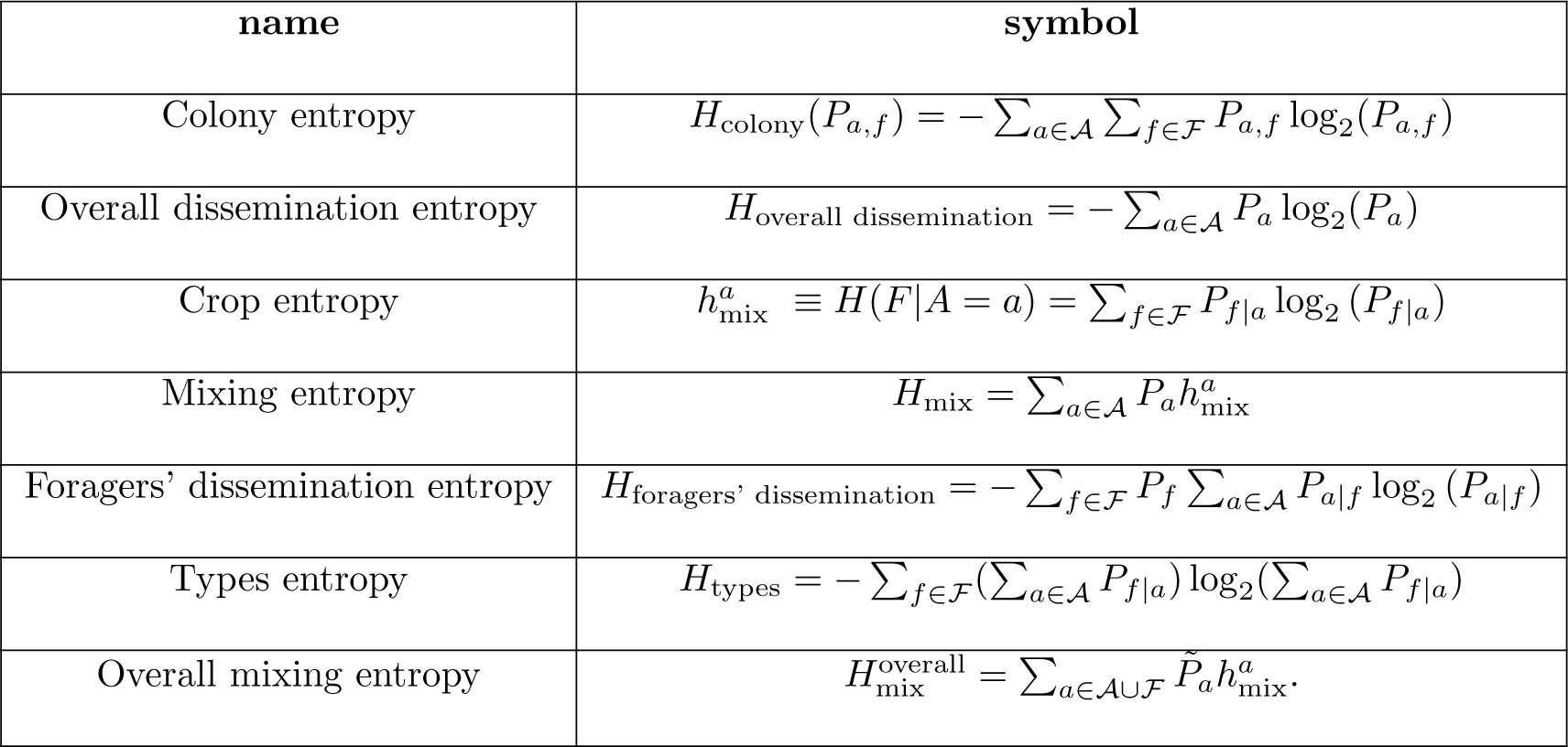
Entropy symbols.

#### Mixing entropy by the largest trophallactic events

Having identified both the global mixing entropy (*H*_mix_) and the food-transfer rule, we aim to reveal how the former emerges from the later. To this end we examine the connection between the volume of trophallactic events and the crop-entropy of individuals 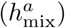 (the mixing entropy is the weighted average of the entropy of the food mixture within the individuals’ crops).

The food-transfer rule suggests that, if not limited by the donor’s food load, in a receiving interaction, an ant is provided with a random volume of food that follows an exponential distribution, with an average that is proportional to her free storage space (*i.e.*, the difference between her capacity and her current load [33]). This implies that, on average, an ant receives food in a series of decreasing volumes. Although this statement is correct only on average, for the estimation of the mixing level the order of the received quantities is not significant

Individual crop entropy can change significantly only if the ant receivers large quantity with respect to the amount she already posses. Therefore, we first calculate the entropy induced by the largest-volume trophallactic events. We find a high correlation between the crop entropy and the entropy calculated for the n-largest events taken from the data of each individual (where *n* is taken as the number of foragers in the experiment a): The distributions of the crop entropy and entropy of the *n* largest interactions are similar (Kolmogorov-Smirnov statistic on 2 samples:KS statistic = 0.15, pvalue= 0.13, Fig. S5a), meaning that the large interactions explain most of the crop entropy.

We further compare the empirical result of crop entropy to a very simple food-accumulation model inspired by the empirical food-transfer rule. Assuming that an ant receives food in a sequence trophallactic interactions, each time of volume that equals to fraction *δ* of her free space, the amount of food received in the j-th interaction equals to:

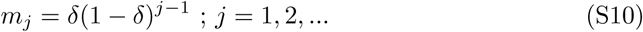

assuming further that each interaction is of a different source type, the entropy of a sequence of n interactions is simply:

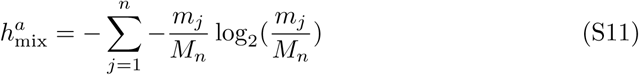

where: 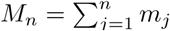

SI Fig. S5 indeed shows that this very simplified accumulation model captures the main trend of the experimental outcome (Kolmogorov-Smirnov statistic on 2 samples: KS statistic = 0.01, pvalue= 0.57.)

## Supplemental Figures

### Supplemental Item

**Movie: Food dissemination in ant colony**. Relates to Fig. 1b-d. Visualization of the process of food dissemination and mixing based on the experimental data. The process begins with foragers, who feed directly at the food source. Food collected by a forager is labeled according to forager identity: forager-268: green, forager-171: orchid, forager-207: red, forager-421: Yellow and forager-180: Blue. Colored blobs overlaid on the movie depict the computationally determined amount of food each ant carries. Each blob is composed of colored transparent layers. The long axis of the ellipse depicting each color is proportional to the amount of food held by the ant which is associated with the forager corresponding to this color (*i.e.*, food that was originally collected at the food source by this forager.).

**Fig. S1.**
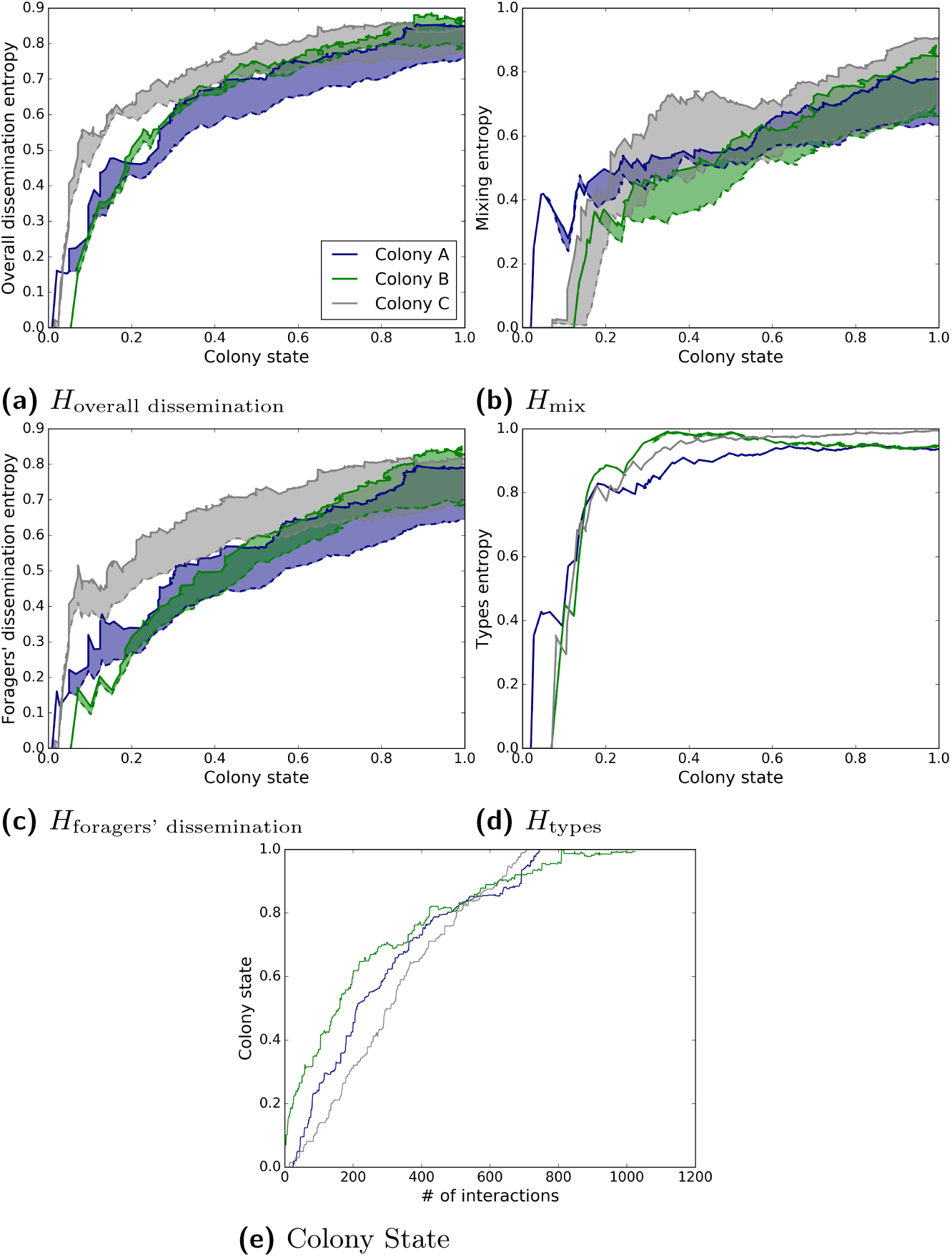
Colony entropy. Related to Fig. 2. Each color stands for a different experiment (blue: colony A (also shown in the main text), green: colony B, grey: colony C). The upper edge of each colored area represents the entropy as calculated from the experimental data. The lower edge (dashed line) depicts the results of a hybrid simulation in which interaction between non-forager workers are excluded. **(a)** Overall dissemination entropy normalized by log(*N*_ants_)). **(b)** Mixing entropy normalized by log(|*ℱ* |)). **(c)** Foragers’ dissemination entropy normalized by log(*N*_ants_)). **(d)** Sources entropy normalized by log(|*ℱ* |)). **(e)** Colony state - normalized total amount of food in the colony as a function of number of interactions.

**Fig. S2.**
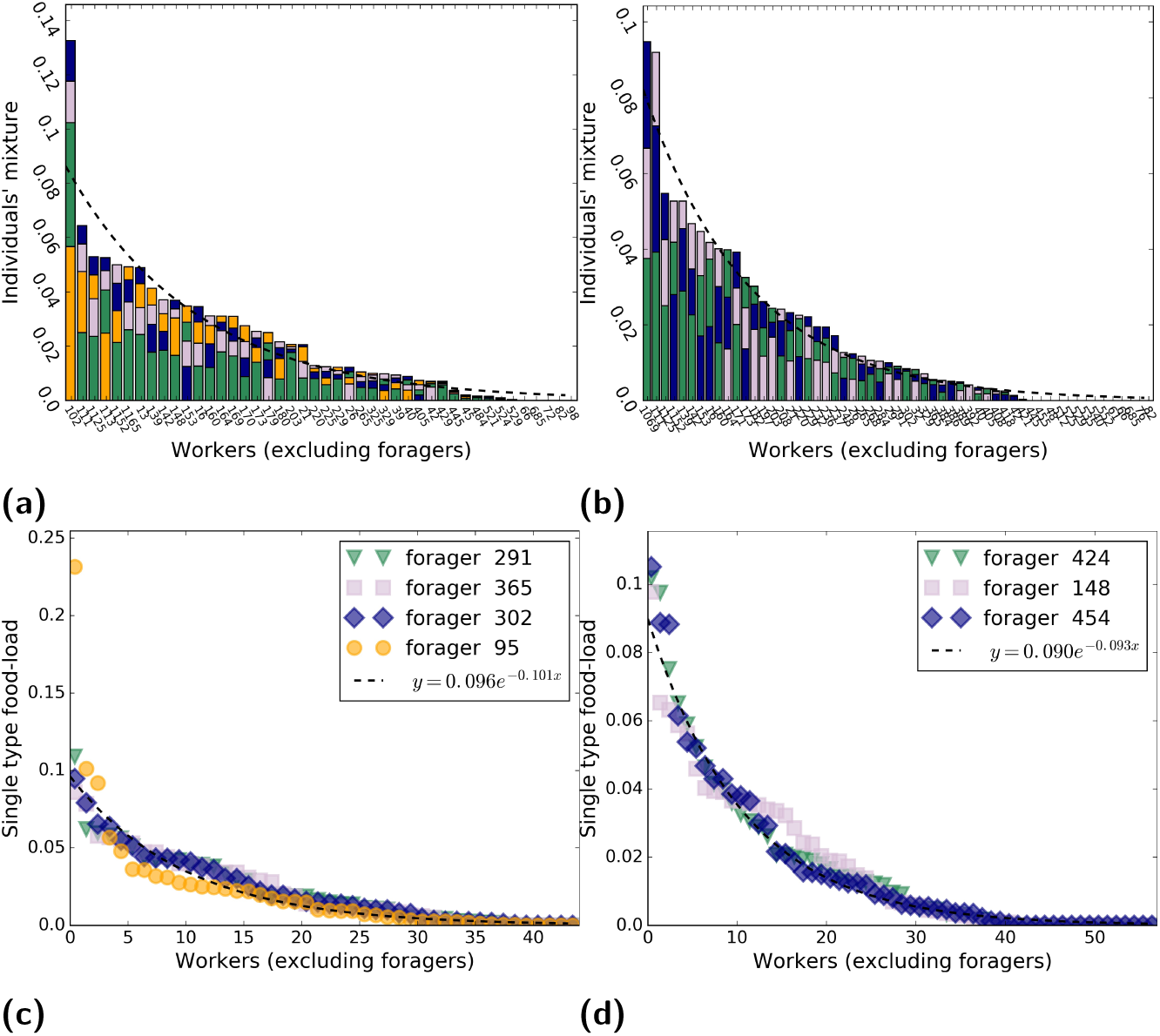
Food distribution. Relates to Fig. 2. **(a and b)** *P* (*a*|*F* = *f*): y-axis represents the fraction of food of source f in ant with index x. Index x (x-axis) is sorted from the largest to the smallest, for colonies B and c respectively. Each color stand for a different forager (*source*). **Dashed line (a)**-fit: *y* = *ae*^*-bx*^, *a* = 0.086 ± 0.005, *b* = 0.089 ± 0.008, *R*^2^ = 0.87.**(b)** Colony C: *P* (*a*|*F* = *f*) **Dashed line (b)**- fit: *y* = *ae*^*-bx*^, *a* = 0.082 ± 0.001, *b* = 0.085 ± 0.002, *R*^2^ = 0.97. **(c and D)** *P* (*f* |*A* = *a*): y-axis represents the fraction of food of source f in ant a. Ants (x-axis) are sorted from the largest to the smallest according to their crop load at the end of the experiment, for colonies B and c respectively. Each color stand for a different forager (*source*) and sorted within each bar according to the fraction of the individual crop load. **Dashed line (c)**- fit: *y* = *ae*^*-bx*^, *a* = 0.09 ± 0.0035, *b* = 0.1 ± 0.0055, *R*^2^ = 0.81. **Dashed line (d)**- fit: *y* = *ae*^*-bx*^, *a* = 0.09 ± 0.0015, *b* = 0.093 ± 0.002, *R*^2^ = 0.93.

**Fig. S3.**
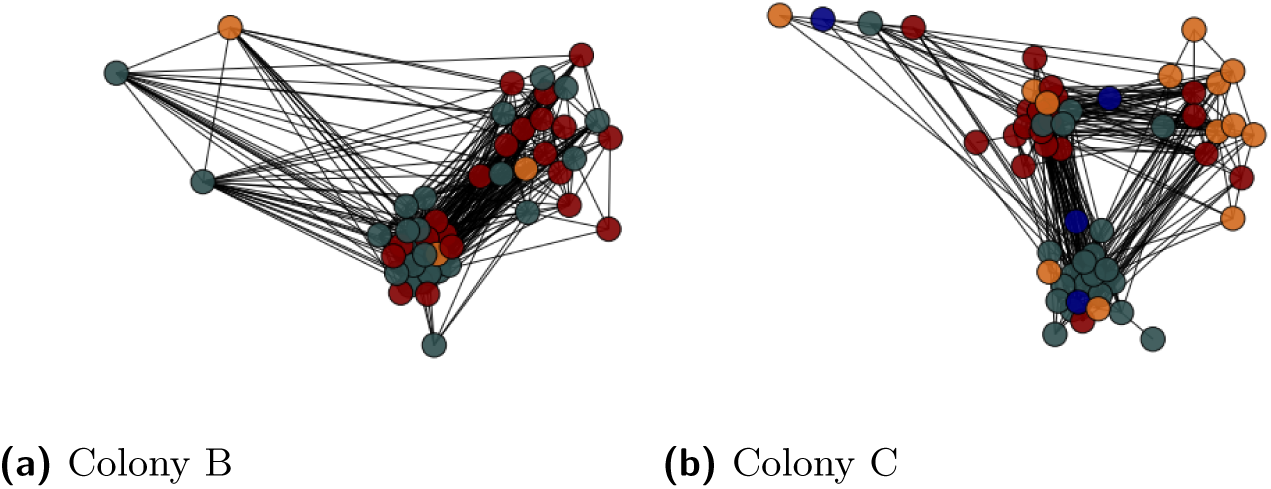
Trophallactic network. Relates to Fig. 3a. Visualization of the undirected trophallactic network in which ants are the vertexes (circles) and interactions are edges (black lines), laid out with the spring embedded layout from Networkx [34] according to communities (colors). (Colony B:) The abundance of inter communities edges is high (177 inter edges and 275 intra edges) and the division to community does not capture the structure of the topology (number of communities=3, transitivity=0.537, modularity=0.128, quality performance=0.65). (Colony C:). This maximal modularity partition shows the same number of intra-community edges (n= 170) as inter-community edges (n = 173) suggesting that division into communities does not capture the n topology of this network (number of communities = 4, transitivity = 0.4, modularity = 0.174, quality performance = 0.7).

**Fig. S4.**
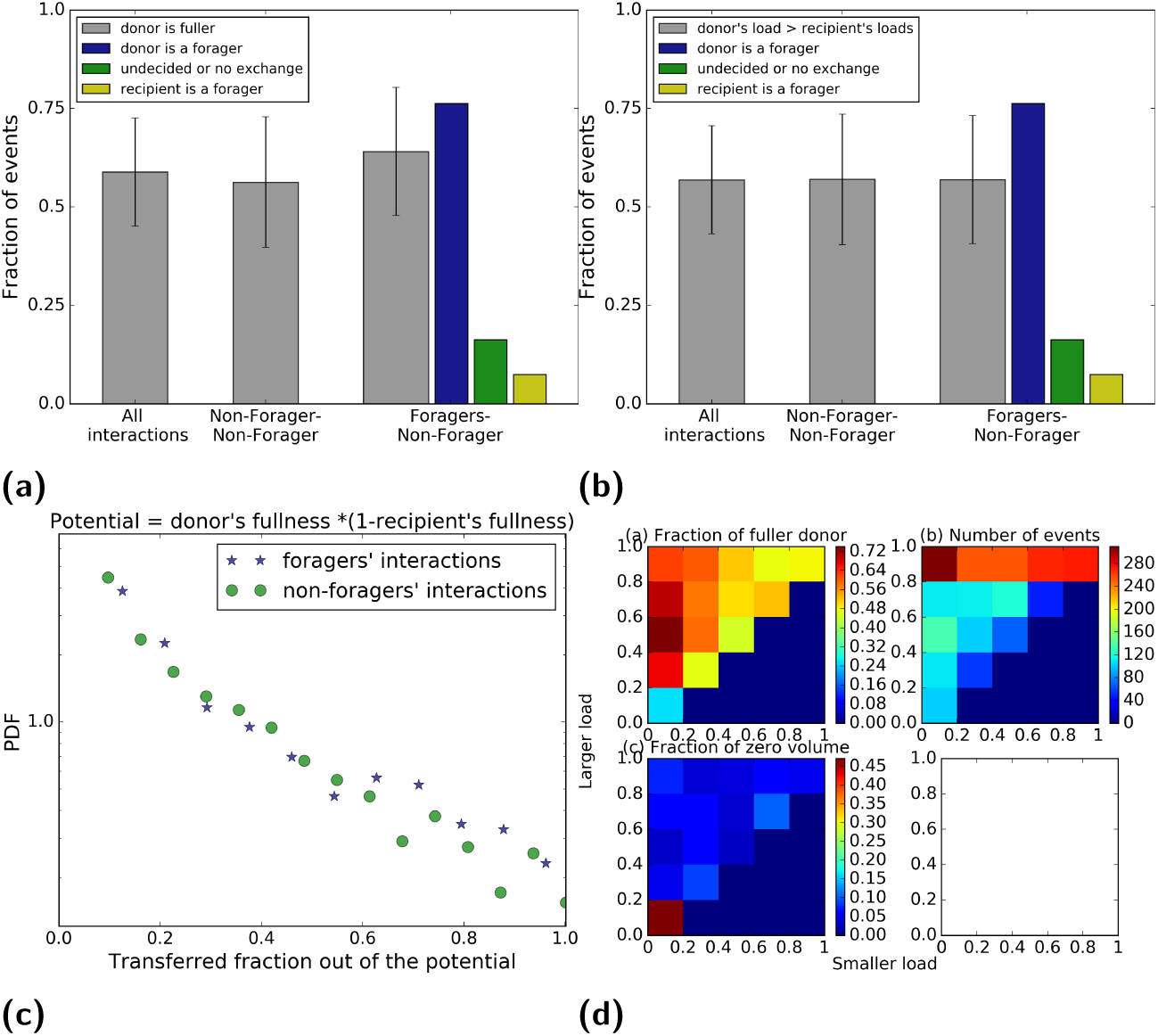
Interaction rules. Relates to Fig. 3b. **(a)** The direction of trophallactic interaction. Grey bars - the fraction of interactions in which the donor was more full for three different subgroups: all-interactions (*N* = 2141, 0.59 ± 0.14), non-forager - non-forager (*N* = 1357,0.56 ± 0.17) forager-non-forager (*N* = 713, 0.64 ± 0.16). Error bar stand for the events in which the transferred volume was below the detection error. The tendency to be higher than 0.5 may be explained by the cases in which the recipient was empty, in this case food can flow in one direction only. Blue bar-fraction of events in which the donor was a forager out of all interactions that include forager and a non-forager worker (*N* = 713,value= 0.76). Green bar-fraction of events in which the direction could not be determine, (either because nothing was transferred or due to measurement error) out of all interactions that include forager and a non-forager worker (*N* = 713,value= 0.16) Yellow bar - fraction of events in which the recipient was a forager out of all interactions that include forager and a non-forager worker-(*N* = 713, value = 0.07). **(b)** Similar to **a** but here the grey bars signify the fraction of interactions in which the donor’ crop load was greater than the recipient’s load: All-interactions (*N* = 2141, 0.59 ± 0.14), non-forager - non-forager (*N* = 1357,0.58 ± 0.17), forager-non-forager (*N* = 713,0.61 + 0.16). **(c)** *δ* rule for foragers and non-foragers: Blue - interaction between foragers and non-forager workers (*N* = 713), Green-interactions between non-forager workers (*N* = 1357). The two cases show no obvious difference (Kolmogorov-Smirnov statistic on 2 samples:KS statistic = 0.067, pvalue= 0.07). **(d) 2d trophallactic direction plots**. Data included all interactions from the three experiments (*N* = 2141), and was binned according to the trophallactic-pair level of satiety as a fraction of the capacity of each ant (x-axis the ant with the lower satiety level, y-axis the ant with the higher satiety level). Colors indicate: **(d-a)** Fraction of interaction in which the donor ant was fuller. **(d-b)** Number of events. **(d-c)** Fraction of near-zero volume events in which direction could not be determined.

**Fig. S5.**
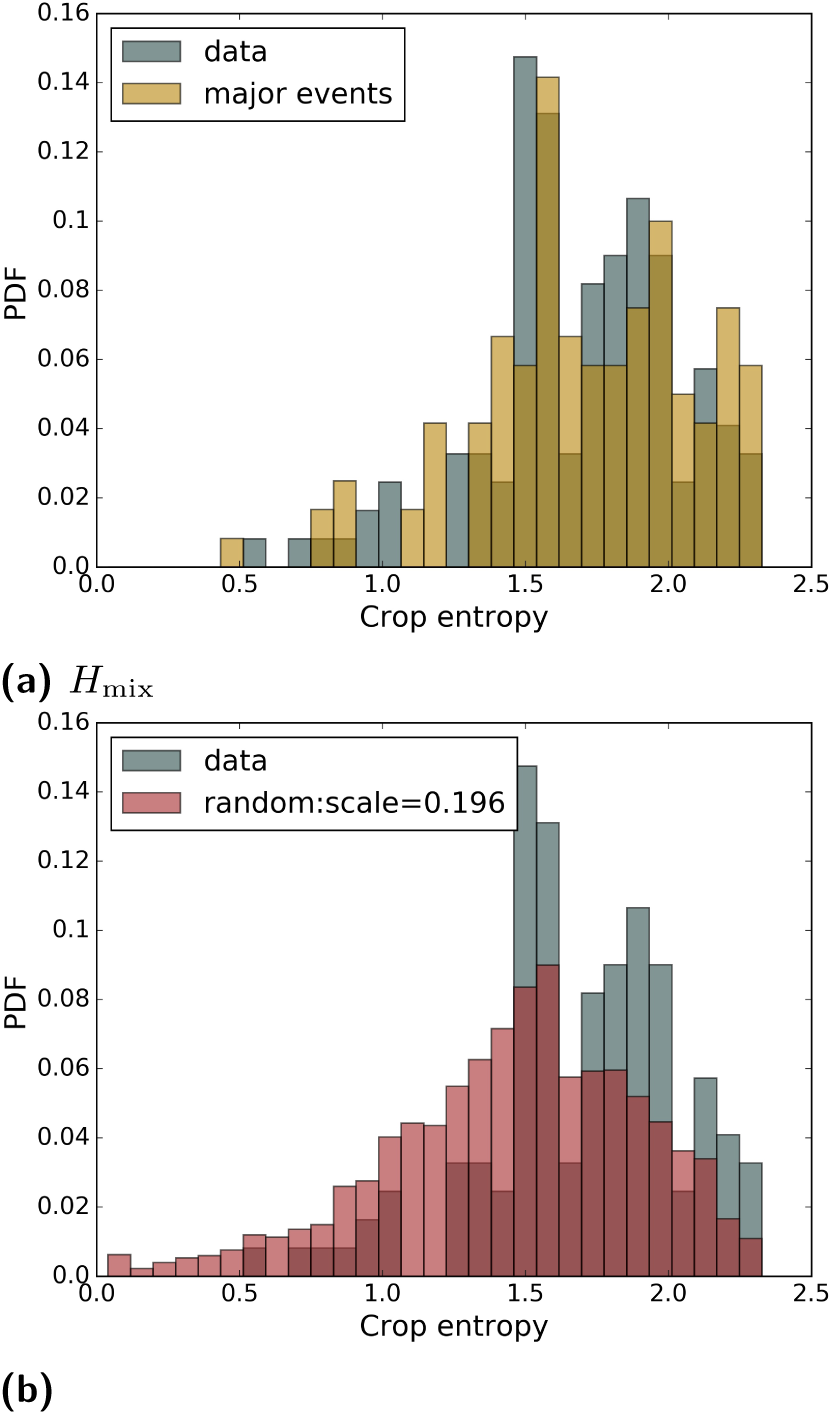
Crop entropy and the entropy generated by the largest events. Relates to Fig. 5 The number of largest events is chosen as the number of foragers in the experiment. **a)** PDF of crop entropy (grey) and the entropy generated by the largest events (yellow). Kolmogorov-Smirnov statistic on 2 samples:KS statistic = 0.15, pvalue= 0.13. **b)** PDF of crop entropy (grey) and the entropy generated by the sequence (red) *m*_1_, *m*_2_,..*M*_*n*_, where *m*_*j*_ = *δ*(1 − *δ*)^*j-*1^, *δ* = ‘scale’ = 0.196 and *n* = number of foragers. Kolmogorov-Smirnov statistic on 2 samples: KS statistic = 0.01, pvalue= 0.57.

## Footnotes

1 Figures numbered S1,… are to be found in the Supplementary Information.

